# Environmental persistence but not per capita transmission rates of a chytrid fungus determines infection prevalence

**DOI:** 10.1101/2021.03.17.435818

**Authors:** Samantha L. Rumschlag, Sadie A. Roth, Taegan A. McMahon, Jason R. Rohr, David J. Civitello

## Abstract

Understanding local-scale variability in disease dynamics can be important for informing strategies for surveillance and management. For example, the amphibian chytrid fungus (*Batrachochytrium dendrobatidis*; Bd), which is implicated in population declines and species extinctions of amphibians, causes spatially variable epizootics and extirpations of its hosts. Outbreak heterogeneity could be driven by differential survival of zoospores, the free-living infectious life stage of Bd, or the persistence of dead zoospores and/or its metabolites in water, which could induce resistance among hosts. To gain a mechanistic understanding of the potential for variation in local transmission dynamics of Bd, we conducted Bd survival and infection experiments and then fit models to discern how Bd mortality, decomposition, and per-capita transmission rate vary among water sources. We found that infection prevalence differed among water sources, which was driven by differences in mortality rates of Bd zoospores, rather than differences in per-capita transmission rates. Specifically, zoospore mortality rates varied significantly among pond water treatments and were lower in artificial spring water compared to pond water sources. These results suggest that variation in Bd infection dynamics could be a function of differences in exposure of hosts to live Bd. In contrast to the persistence of live zoospores, we found that rates of decomposition of dead zoospores did not vary among water sources. These results may suggest that exposure of hosts to dead Bd or its metabolites, which have been shown to induce acquired resistance, might not commonly vary among nearby sites. Ultimately, a mechanistic understanding of the drivers of variable epizootics of Bd could lead to increases in the effectiveness of surveillance and management strategies.

## Introduction

Strategies for the surveillance and management of wildlife infectious diseases are informed by a mechanistic understanding of the distribution and persistence of pathogens in host populations and the environment (Cunningham et al. 2017, Barnett and Civitello 2020). Yet, characterizing these mechanisms remains immensely challenging because of the complexity and natural variability of host-pathogen systems and the environments in which they exist (Plowright et al. 2008, Tompkins et al. 2011, Gandon et al. 2016). For instance, infection prevalence can vary widely among neighboring host populations (Ostfeld et al. 2005, Favier et al. 2005, Wood et al. 2007, Munster et al. 2007). Some of this local variability may be a function of the net effects of environmental conditions on host exposure to pathogens and host susceptibility to infection (Rohr et al. 2008a, Civitello et al. 2013, Civitello and Rohr 2014, Rumschlag et al. 2019). Spatial variation in environmental conditions may influence mortality of infectious, free-living pathogens, leading to differences in host exposure. For instance, the survival of free-living stages of many marine trematode species are sensitive to temperature, salinity, and pH which leads to spatial variability in their abundance in intertidal zones (Koprivnikar et al. 2010, Lei and Poulin 2011). At the same time, environmental variability could lead to differences in transmission of pathogens through effects on host susceptibility to infections. For instance, the herbicide atrazine can increase susceptibility of amphibian hosts to infection with a trematode parasite by compromising host immunity, leading to variability in infection loads across space (Rohr et al. 2008b). The net effects of environmental conditions on host exposure to pathogens and host susceptibility to infection may explain spatial variability in the prevalence of pathogens (Tompkins et al. 2011, Estrada-Peña et al. 2014). For many wildlife disease systems, the influence of local environmental conditions on exposure and per capita transmission has not been adequately evaluated, which is likely leading to challenges in the development of effective strategies for disease surveillance and management.

Understanding how local environmental variability impacts host-pathogen systems can be particularly important when associated with catastrophic host declines. *Batrachochytrium dendrobatidis* (Bd), for example, is a generalist pathogen that causes the disease chytridiomycosis in amphibians and has been implicated in population declines, extirpation and extinctions of hundreds of amphibian species worldwide (Kilpatrick et al. 2010, Venesky et al. 2014b). Bd is an aquatic pathogen with a motile and infectious zoospore stage, which penetrates and colonizes host tissues containing keratin (Berger et al. 1998, McMahon et al. 2013a, McMahon and Rohr 2015). After colonization, zoospores develop into zoosporangia, which produce and release new zoospores. New zoospores re-infect the same host or are released into the environment (Berger et al. 2005).

Host population persistence and response to Bd are heterogenous (Brannelly et al. 2021); some amphibian host populations are suddenly extirpated, while others persist with endemic Bd transmission (Pilliod et al. 2010, Briggs et al. 2010, Longo and Burrowes 2010). While many studies have investigated sources of variability in Bd infection patterns at large spatial scales (regional to global), which include temperature regime (Liu et al. 2013, James et al. 2015, Cohen et al. 2017), precipitation (Becker and Zamudio 2011, James et al. 2015), altitude (Muths et al. 2003, Rohr and Raffel 2010, Lambertini et al. 2021), host community diversity (Venesky et al. 2014a, Cohen et al. 2016), and land use practices (Padgett-Flohr and Hopkins 2010, Rumschlag and Rohr 2018, Rumschlag and Boone 2020), less is known about the mechanisms that drive local transmission dynamics. For instance, currently, the relative importance of environmental variation in driving Bd exposure and per capita transmission remains uncertain, which could have important ramifications for explaining patterns of Bd infection prevalence across host populations.

Neighboring populations of amphibian hosts of the same species can vary tremendously in their Bd infection prevalence (Raffel et al. 2010, Strauss and Smith 2013, Venesky et al. 2014a, Chestnut et al. 2014, Rumschlag and Boone 2020). This local-scale variability in patterns of infections could be caused by variation in zoospore mortality, leading to differences in Bd exposure among sites, or by variation in host susceptibility to Bd infection. The main environmental sources of Bd mortality are likely consumption by predators and poor water quality. Protozoa, zooplankton, and filter feeding tadpoles can consume zoospores suspended in the water column, decreasing Bd exposure, and lowering infection rates and loads in amphibian hosts (Buck et al. 2011, Hamilton et al. 2012, Searle et al. 2013, Venesky et al. 2014a, Schmeller et al. 2014). In addition, water quality parameters including pH (Piotrowski et al. 2004, Kärvemo et al. 2018), salinity (Heard et al. 2014, Stockwell et al. 2015, Clulow et al. 2018), temperature (Voyles et al. 2012, Raffel et al. 2015), and the presence of contaminants (McMahon et al. 2013b, Rumschlag et al. 2014, Rohr et al. 2017, Rumschlag and Rohr 2018) can influence the survival of zoospores and ability to infect hosts. Simultaneously, local scale variability in infection prevalence could be a result of differences in host susceptibility. In particular, susceptibility of species may vary as a result of differences in water quality (Jani and Briggs 2018, Greenspan et al. 2019) or intraspecific population-level differences in susceptibility (Bradley et al. 2015).

In addition, dead Bd zoospores and/or the metabolites they produced can induce acquired resistance in amphibian hosts (Ellison et al. 2014, McMahon et al. 2014, Savage et al. 2016). If environmental conditions cause zoospores to die rapidly once they are released from zoosporangia, hosts could be exposed regularly to dead Bd zoospores. However, if after death Bd decomposes quickly, host exposure to dead Bd zoospores could be minimal. Spatial variability in acquired resistance of hosts through variability in the persistence of dead Bd and its metabolites could contribute to spatial variability in Bd outbreaks. If composition and densities of zoospore decomposers and scavengers and water quality influence of the persistence of dead Bd or its metabolites in the environment, then variation in these factors at local spatial scales could lead to variation in Bd disease dynamics.

To gain a mechanistic understanding of the potential for variation in local transmission dynamics of Bd, we conducted experiments and fit transmission models to discern how Bd mortality, decomposition, and per capita transmission vary among water sources. We predicted that rates of Bd zoospore mortality and decomposition would be higher in pond water than artificial spring water (ASW) because ASW lacks any zoospore predators, competitors, and scavengers. High zoospore mortality, in turn, should lead to lower rates of transmission and infection prevalence in frog hosts in pond water compared to ASW. We also hypothesized that there would be significant variation among pond water treatments in rates of Bd zoospore mortality, decomposition, and infectivity, perhaps because of differences in abiotic conditions or zoospore predators, competitors, and scavengers that can affect environmental persistence of Bd.

## Materials and Methods

### Animal Collection and Care

We collected multiple clutches of recently hatched Cuban treefrog (*Osteopilus septentrionalis*) tadpoles at Gosner stage 25 (Gosner 1960) from University of South Florida Botanical Gardens. We maintained the tadpoles in the laboratory for nine days until the start of the second experiment (described below) at 21 ± 1°C with a 14:10 hr light:dark photoperiod in tanks containing 90% artificial spring water (ASW; Cohen et al. 1980) and 10% rainwater. Tadpoles were fed *ad libitum* a mixture of fish flakes and spirulina suspended in agar, and water was changed daily. Tadpole survival was checked daily, and no mortality was observed prior to the start of the second experiment.

### Bd Inoculate Preparation

For both experiments, Bd (isolate SRS 810; isolated from *Lithobates catesbeianus* in USA) was grown on 1% tryptone agar plates at 23°C over 7-10 d. Immediately before use in each experiment, Bd plates were flooded with DI water for 30 min to suspend zoospores. Water containing zoospores was homogenized among plates. The zoospore concentration was determined using a hemocytometer and the solution was diluted to the given target concentration (described below).

### Experimental Designs

To quantify rates of Bd mortality, decomposition, and per capita transmission in pond water and ASW, we conducted two experiments and used the results of the experiment to inform models estimating Bd mortality, decomposition, and per capita transmission. In the first experiment, we measured concentration of live and dead Bd zoospores in four water treatments (ASW or pond water from one of three locations: Flatwoods Park, Green Swamp Wilderness Preserve, or Trout Creek Conservation Park) by destructively sampling experimental units across time (0, 6, 16, 24, or 48 hrs). An experimental unit was 1 ml of zoospores at a concentration of 125,000 zoospores/ml plus 1 ml of a given water treatment, and each water treatment-time combination was replicated 10 times for a total of 200 experimental units. These three sources of pond were selected because previous studies surveying amphibian populations for Bd infections at these three locations have not detected Bd (*unpublished data*). Pond water was collected from the end of the ponds and was not filtered because we wanted to maintain the natural zoospore predators, competitors, and scavengers. ASW was used to control for the presence of any zoospore predators, competitors, and scavengers.

Survival of zoospores was measured via counts of live and dead zoospores that were stained with Trypan blue (McMahon and Rohr 2014). We added equal parts Trypan and zoospore-water treatment solution before counting live and dead zoospores on a hemocytometer. Cell counting of samples was randomly and evenly assigned to two people. Dissolved oxygen and pH were taken from the stock sample for each of the water treatments used in the experiment: ASW (9.20 mg/L DO, 6.74 pH), Flatwoods (8.50 mg/L DO, 6.66 pH), Green Swamp (9.15 mg/L DO, 7.30 pH), and Trout Creek (9.40 mg/L DO, 6.38 pH). All water treatments were maintained 21 °C during the experiment.

In a second experiment, we exposed Cuban treefrog tadpoles (*Osteopilus septentrionalis*) to Bd zoospores in four water treatments (ASW or pond water from one of three locations: Flatwoods Park, Green Swamp Wilderness Preserve, or Trout Creek Conservation Park) at five time lags between the introduction of Bd live zoospores (5×10^5^ zoospores/ml) and the introduction of the tadpole (0, 3, 6, 12, and 24 hrs) in 100 ml of the given water treatment. Each water treatment-time lag combination was replicated ten times with a single tadpole as the experimental unit for a total of 200 tadpoles. Tadpoles were exposed to Bd in their given water and time lag treatments for 24 hrs. Then, every tadpole was transferred to 450 ml ASW water for two weeks so that they could develop detectable infection, if present. There was no tadpole mortality. Dissolved oxygen and pH were taken from the stock sample for each of the water treatments used the second experiment: ASW (7.0 mg/L DO, 7.52 pH), Flatwoods (7.3 mg/L DO, 6.76 pH), Green Swamp (7.9 mg/L DO, 7.84 pH), and Trout Creek (6.5 mg/L DO, 6.98 pH). The experiment was conducted at 21 ± 1°C with a 14:10 hr light:dark photoperiod. During the experiment, tadpole water was changed once per week and tadpoles were fed *ad libitum* a mixture of fish flakes and spirulina suspended in agar.

At the end of two weeks, tadpoles were euthanized, and their mouthparts were excised and analyzed for Bd using quantitative PCR (qPCR). To determine the Bd quantity on each tadpole, we followed the qPCR protocol described by Hyatt et al. (2007). Briefly, we used PrepMan Ultra (Applied Biosystems) to extract DNA. To increase extraction efficiency, the mouthpart tissue was placed in a cell disruptor (Disruptor Genie, Scientific Industries) and agitated with 0.035 ± 0.005 g of zirconia/silica beads for a total of 2.25 minutes. We used TaqMan Exogenous Internal Positive Control Reagents (Applied Biosystems) to screen for inhibition in all samples. We diluted and re-ran any samples with inhibition.

### Mortality, decomposition, and per capita transmission modeling and analysis

We estimated the rates of mortality, decomposition, and per capita transmission of Bd zoospores by fitting a model to the results of the survival and infection experiments. The model used coupled ordinary differential equations to track the concentrations of live zoospores, *Z_L_*, dead zoospores, *Z_D_*, susceptible frogs, *S*, and infected frogs, *I* (Eqs. 1-4, Fig. 1). It assumes a constant background mortality of live zoospores, at per capita rate *m*, a constant decomposition rate of dead zoospores, at per capita rate *d*, and density-dependent transmission via zoospore-frog contact, at transmission rate β. For tractability, it assumes no host mortality and that depletion of zoospores via infection (*sensu* Civitello and Rohr 2014) is negligible relative to zoospore mortality:

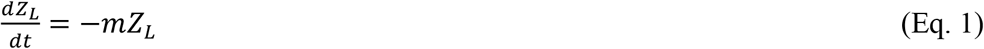

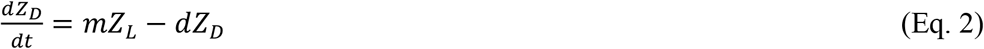

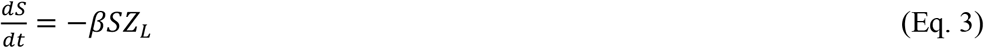

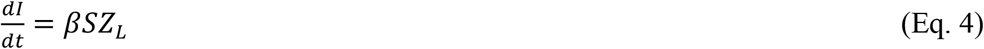

**Figure 1.**
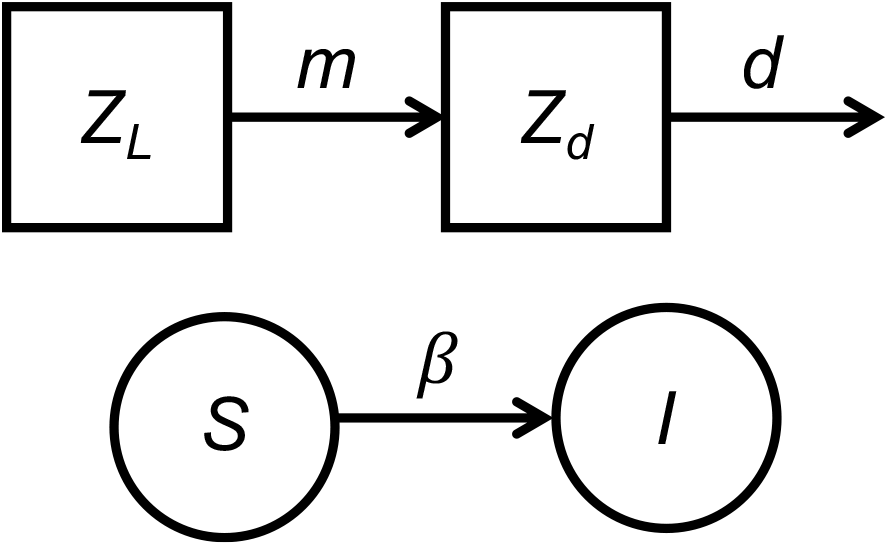
Conceptual diagram of the model which uses coupled ordinary differential equations to track the concentrations of live zoospores, *Z_L_*, dead zoospores, *Z_D_*, susceptible frogs, *S*, and infected frogs, *I*. The model assumes a constant background mortality of live zoospores, at per capita rate *m*, a constant decomposition rate of dead zoospores, at per capita rate *d*, and density-dependent transmission via zoospore-frog contact, at transmission rate β.

Integrating this model yields analytical solutions for the key quantities observed in the experiments: *Z_L_*(t), *Z_D_*(t), and *S*(t). Specifically, *Z_L_*(t) follows an exponential decay function, depending on the mortality rate, *m*, the initial density of live zoospores, *Z_L_*(0), and time, *t*:

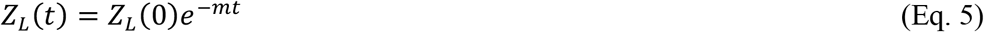

Assuming that there are initially no dead zoospores, *Z_D_*(0) = 0, yields an analytical solution for *Z_D_*(t):

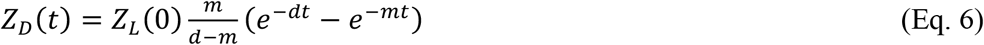

These equations predict the density of live and dead zoospores observable at time *t*. We fit these functions to paired observations of these quantities from the zoospore survival experiment. We assumed independent log-normal errors to obtain likelihoods:

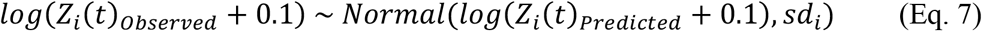

where *i* = L, D and *sd_i_* is the standard deviation of predictive errors. We added a small quantity, 0.1, to each observation and prediction to avoid taking log(0).

The model also yields an analytical solution for *S*(t), which provides a well-established method for estimating the per capita transmission rate, β, via a likelihood function derived from the binomial distribution:

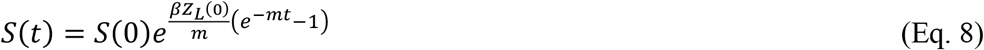

The above equation assumes that hosts and parasites are placed in the environment together at time *t* = 0. However, in our infection experiments, we added zoospores to containers at time *t* = 0, and then added a single susceptible host after a time lag, *t*, such that *S*(*t*) = 1. We then removed the host individual after an exposure period of length τ (in our experimental τ = 1 day for all treatments). Thus, the tadpole is predicted to be exposed upon its introduction to the concentration of zoospores remaining alive at time *t*, *Z_L_*(*t*), and the entire trial ends at the total time *t* + τ. Incorporating these experimental conditions into Eq. 8 yields a specific prediction for *S*(*t +* τ):

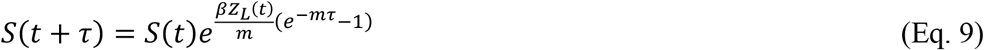

Substituting *Z_L_*(*t*) from Eq. 5 and simplifying yields:

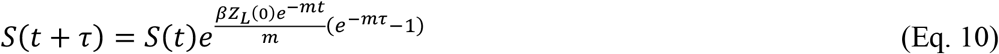

Because the total density of tadpoles never changes, *S*(*t*) and *S*(*t* + τ) fully specify prevalence at time *t* + τ, *p*(*t* + τ), which we use in the binomial likelihood function:

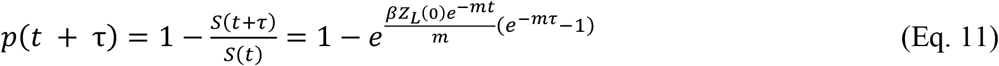

This quantity specifies the probability of infection in the binomial likelihood, which we calculate for each tadpole, which is diagnosed as infected or uninfected (Rachowicz and Briggs 2007). We combined these likelihood functions (Equations 7 and 11) to generate an overall likelihood for all observations (live zoospores, dead zoospores, and infected hosts) across the two experiments.

With the theoretical and statistical models in place, we estimated parameters (zoospore mortality, zoospore decomposition, and per capita transmission) for a series of 15 transmission models (see Table 1). We tested models that assumed that these three parameters were either constant across all water sources, could vary across all four sources, or could vary between ASW versus the pond waters (as a group). We evaluated the relative performance of this set of 15 models using AICc, Akaike model weights, and importance scores (Burnham and Anderson 2002).

**Table 1.** Results of the model competition analysis for 15 models fit to the Bd zoospore mortality, decomposition, and infection data. Models differed based on whether mortality (*m*), decomposition (*d*), and transmission (β) parameters were fit with a single, common estimate among the four water sources (Parameters with Single Estimates), with two estimates that corresponded to whether the water source was ASW or pond water (Parameters with Two Estimates), or with four estimates that corresponded to each water source (ASW or pond water from Flatwoods Park, Green Swamp Wilderness Preserve, or Trout Creek Conservation Park). Model competition results suggest that mortality, *m*, varied among water sources, but the other two parameters do not.

| Parameters with Single Estimates | Parameters with Two Estimates | Parameters with Four Estimates | df | AICc | ΔAICc | weight |
| --- | --- | --- | --- | --- | --- | --- |
| $\beta$ | - | $m, d$ | 12 | 1377.9 | 0 | 0.40 |
| - | - | $m, d, \beta$ | 15 | 1379 | 1.1 | 0.24 |
| $d, \beta$ | - | $m$ | 9 | 1379.2 | 1.2 | 0.22 |
| $d$ | - | $m, \beta$ | 12 | 1380.2 | 2.2 | 0.13 |
| $\beta$ | $m, d$ | - | 8 | 1385 | 7.1 | 0.01 |
| $d, \beta$ | $m$ | - | 7 | 1389.1 | 11.2 | 0.002 |
| | $m, d, \beta$ | - | 9 | 1389.7 | 11.8 | 0.001 |
| $m, \beta$ | $d$ | - | 7 | 1391.8 | 13.9 | <0.001 |
| $m$ | $d, \beta$ | - | 8 | 1392 | 14.1 | <0.001 |
| $d$ | $m, \beta$ | - | 8 | 1393.9 | 15.9 | <0.001 |
| $m, \beta$ | - | $d$ | 9 | 1395 | 17.1 | <0.001 |
| $m, d, \beta$ | - | - | 6 | 1396.5 | 18.6 | <0.001 |
| $m, d$ | $\beta$ | - | 7 | 1396.7 | 18.8 | <0.001 |
| $m$ | - | $d, \beta$ | 12 | 1397.2 | 19.3 | <0.001 |
| $m, d$ | - | $\beta$ | 9 | 1398.7 | 20.8 | <0.001 |

## Results

The competition of the 15 transmission models indicated that the model with mortality rate, *m*, varying among water sources and a common estimate for decomposition, *d*, and per capita transmission, β, rates fit the data from our zoospore persistence and infectivity experiments best (Table 1). Three models had ΔAICc scores less than two, all of which contained varying parameter estimates for *m* according to the four water sources. None of the models that included varying estimates of *d* and β were better (i.e., more than two AICc lower) than the model that varied *m* alone. In corroboration, mortality rate fit with four estimates received an importance score of almost one, and per capita transmission rate fit with a single estimate received a greater importance score than per capita transmission rates fit with two or four estimates (Fig. 2A). While the importance score for decomposition rate was highest when it was fit with four estimates (Fig. 2A), this result likely reflected a small difference between decomposition in ASW and pond water from Trout Creek (Fig. 2B). A log-likelihood ratio test supported that the lowest AICc model with the additional complexity of *m* and *d* fit with four parameters was not a significant improvement above the model with a single parameter estimate for *d* and β and four parameter estimates according to water treatments for *m* (χ^2^ = 7.581, *df* = 3, *p* = 0.056). So, we have strong evidence that *m* varied among water sources and weak evidence that *d* and β varied among water sources.

**Figure 2.**
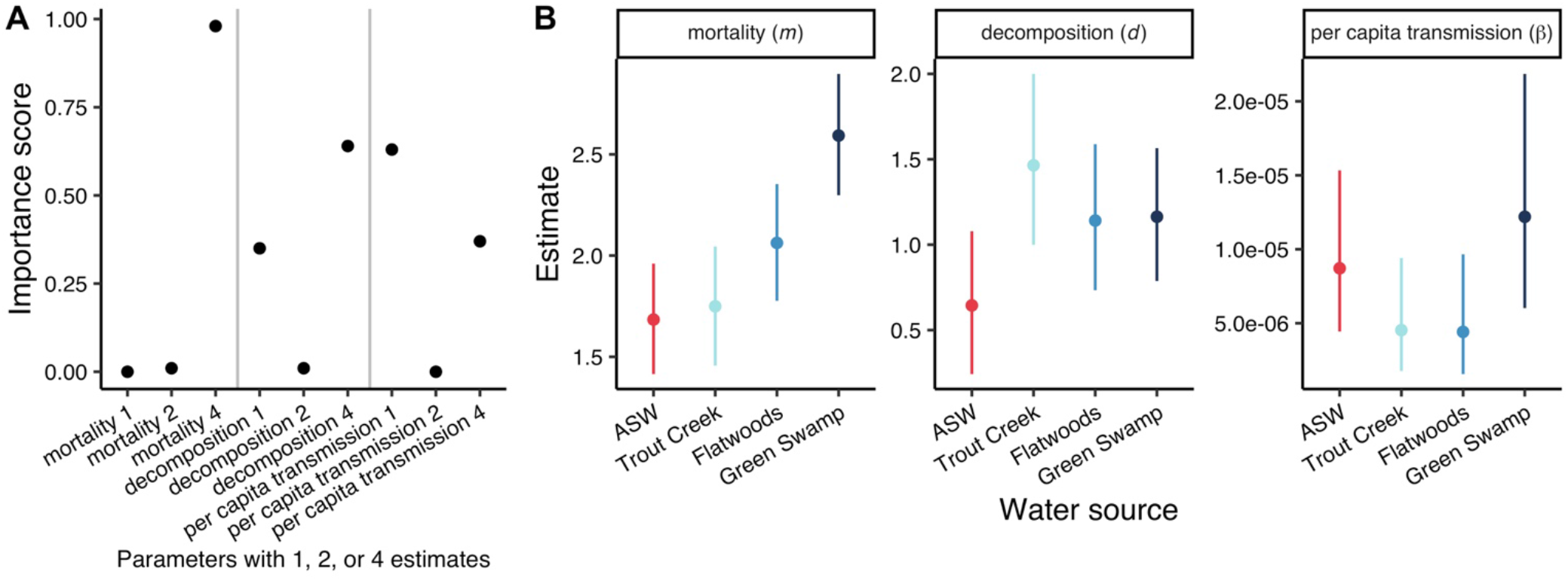
**A)** Importance scores of mortality, *m*, decomposition, *d*, and per capita transmission, β, parameters fit with single estimates, two estimates (ASW or pond water), or four estimates (ASW or pond water from Flatwoods Park, Green Swamp Wilderness Preserve, or Trout Creek Conservation Park). Importance scores were calculated as the sum of the Akaike weights of models including the given parameter (Table 1). Importance scores were greatest for mortality and decomposition rates fit with four estimates according to water sources, while importance score was greatest for per capita transmission fit with a single estimate. **B)** Parameter estimates for mortality, *m*, decomposition, *d*, and per capita transmission, β, from a model including four separate estimates according to water source for each parameter. Error estimates are 95% confidence intervals. Mortality varied significantly among water treatments with the lowest rates of mortality in ASW. Decomposition did not show as great of differences among water treatments compared to mortality, though the largest differences in decomposition were between ASW and pond water from Trout Creek. Per capita transmission did not vary among the four water sources.

Model predictions supported lower rates of zoospore mortality in ASW compared to pond water (Fig. 2B). The expected lifespan of a zoospore (± 95% CI) was 14.4 ± 2.3 hrs in ASW, 13.3 ± 2.2 hrs in pond water from Trout Creek, 11.4 ± 1.6 hrs in pond water from Flatwoods, and 9.4 ± 1.1 hrs in pond water from Green Swamp. There were no obvious patterns between mortality rate of zoospores and DO and pH measurements across water treatments. These differences in mortality rates led to differences in the concentrations of live and dead zoospores across time (Fig. 3). The model predicted that the concentration of live zoospores in ASW was consistently greater across time than the concentration of live zoospores in water from any of the three ponds (Fig. 3A).

**Figure 3.**
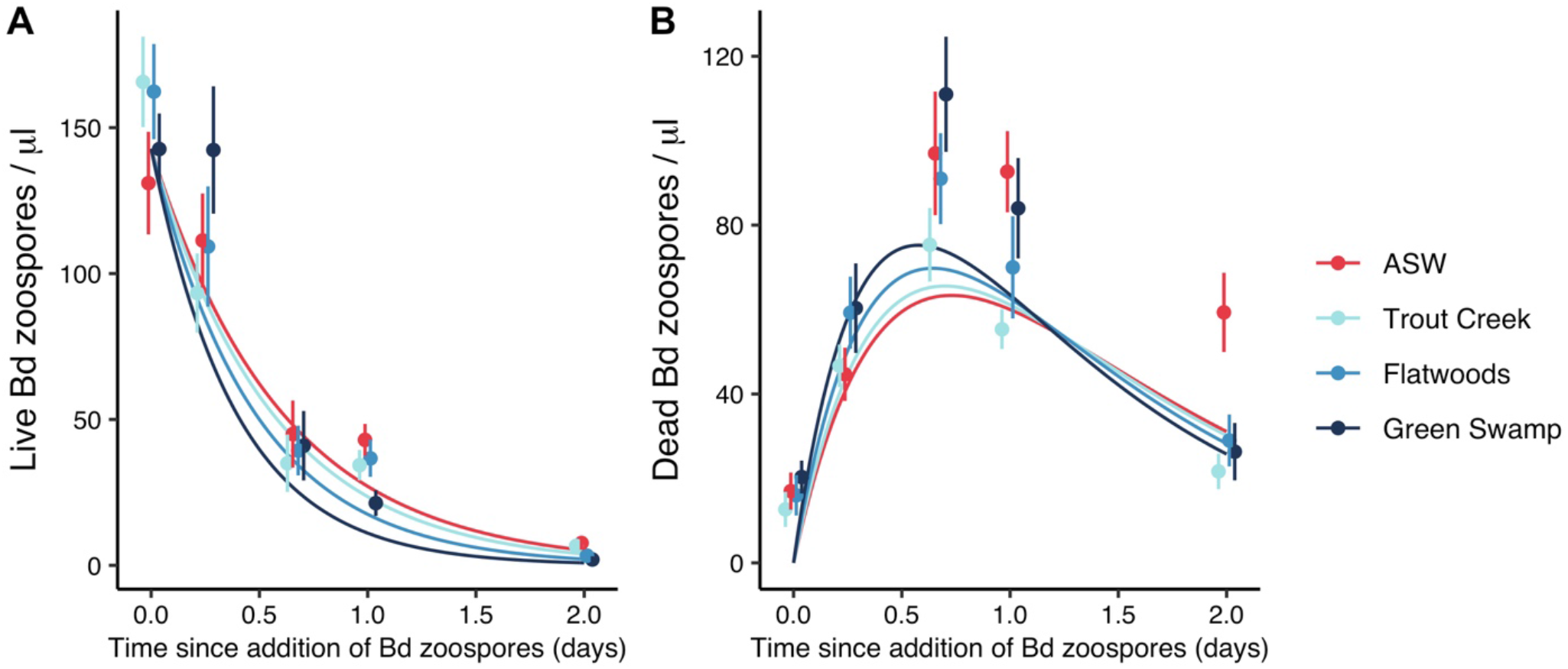
Best-fit predictions of concentrations of **A)** live and **B)** dead zoospores across time from the transmission model. **A)** The concentration of live zoospores decreased across time, driven by differences in mortality rates across the four water treatments (ASW or pond water from Flatwoods Park, Green Swamp Wilderness Preserve, or Trout Creek Conservation Park). The highest concentrations of live zoospores were in ASW, as predicted. Measured means and standard errors of concentrations of live zoospores across time are provided. **B)** The concentration of dead Bd zoospores over time was non-monotonic, with an initial increase as zoospores died, driven by differences in mortality rates across water sources, and then a decrease after 16 hrs, caused by decomposition or scavenging of dead zoospores. The model predicted the concentration of dead zoospores did not differ across water treatments; although at 48 hrs, the measured concentration of dead zoospores was significantly greater in ASW, suggesting lower rates of decomposition or scavenging in accordance with predictions. Measured means and standard errors of concentrations of dead zoospores across time are provided.

While the zoospore mortality rates varied across water treatments, the best model suggested that decomposition rates of zoospores did not. Overall, the expected persistence time of a zoospore (± 95% CI) was 21.6 ± 5.2 hrs. Persistence time in the model was defined as the average time from the death of a zoospore to complete decomposition. The concentration of dead Bd zoospores over time was non-monotonic, with an initial increase as zoospores died and then a decrease after 16 hrs, caused by decomposition of dead zoospores (Fig. 3B). While the rate of decomposition of zoospores did not vary across water treatments, the mean concentration of dead zoospores experimentally measured was substantially greater in ASW at the 48 hr experimental time point (Fig. 3B), suggesting lower rates of decomposition in accordance with predictions.

The best model suggested that the inferred per capita rate of transmission, β, was 7.1 × 10^-6^ ± 2.6×10^-6^ (95% CI), and that this rate did not vary across water treatments. Predicted infection prevalence, which depends on both per capita rate of transmission, β, and zoospore mortality rate, *m*, decreased across time as zoospores died (Fig. 4A). Differences in zoospore mortality rate, but not per capita transmission rate, led to differences in infection prevalence across time; infection prevalence was greater in ASW compared to pond water sources across time (Fig. 4A). The transmission model also predicted a positive association between host infection prevalence and concentration of live zoospores with the highest rates of infection prevalence and the greatest concentration of live zoospores occurring in ASW (Fig. 4B).

**Figure 4.**
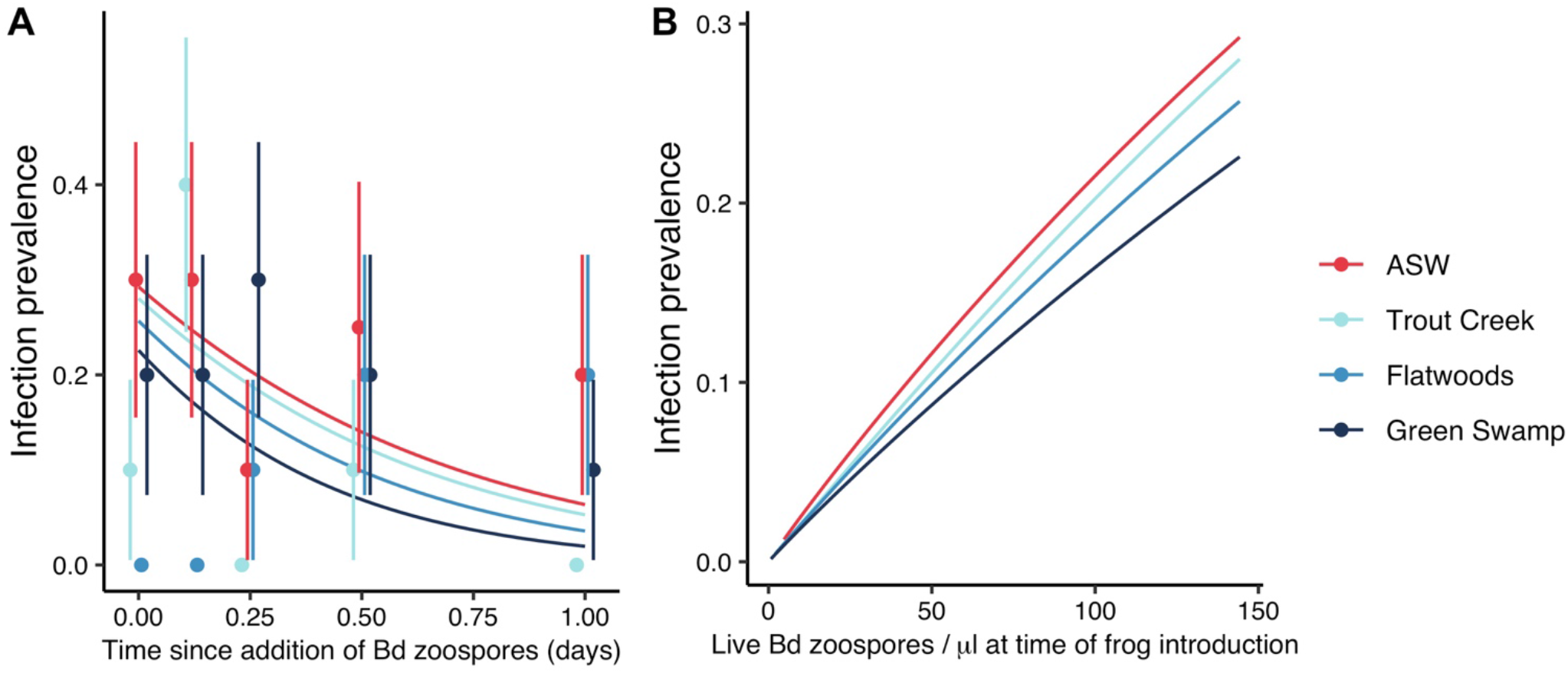
**A)** Best-fit predictions for prevalence of Bd infections across time from the transmission model. Infection prevalence, which was informed by both zoospore mortality and transmission, decreased across time as zoospores died. Differences across water sources in zoospore mortality, but not per capita transmission, led to differences in infection prevalence with the highest infection prevalence across time in ASW. Measured means and binomial standard errors of infection prevalence are provided. **B)** Best-fit predictions for prevalence of Bd infections against concentration of live Bd zoospores. The transmission model predicts a positive association between host infection prevalence and concentration of live zoospores with the highest rates of infection prevalence and the greatest concentration of live zoospores occurring in ASW.

## Discussion

To gain a mechanistic understanding of the potential for local variation in transmission dynamics of Bd, we conducted Bd survival and infection experiments and then fit models to evaluate how Bd mortality, decomposition, and per-capita transmission rates varied among water sources. Results showed that infection prevalence differed among water sources, which was driven indirectly by differences in mortality rates of Bd zoospores and not by differences in per-capita transmission rates. More specifically, zoospore mortality rates varied among pond water sources and were lower in ASW compared to pond water. These differences among pond water sources suggest that local variation in Bd infection dynamics could be a function of differences in Bd exposure. In contrast to the differences in mortality rates of live zoospores among water sources, we found that rates of decomposition of dead zoospores did not vary. These results could suggest that exposure of amphibian hosts to dead Bd and its metabolites, which have been shown to induce acquired resistance (Ellison et al. 2014, McMahon et al. 2014, Savage et al. 2016), might not commonly vary among nearby sites.

Consistent with our predictions, we found that Bd mortality rates varied among pond water sources and that the lowest rates of mortality occurred in ASW compared to pond water. We hypothesized that differences in the predators and/or competitors of live Bd zoospores drove the differences between ASW and pond water. Protozoa and zooplankton consume live zoospores in the water column, decreasing live zoospore persistence and thus infection rates and loads in amphibian hosts (Buck et al. 2011, Hamilton et al. 2012, Searle et al. 2013, Schmeller et al. 2014). Among ponds, mortality rates differed (e.g. Trout Creek versus Green Swamp, Fig. 2B), which conceivably could have been driven by differences in the densities and/or compositions of predators and consumers. While trends in mortality rates across water treatments did not align with pH and dissolved oxygen measurements, given the relatively low sample size, the direct effects of water quality on Bd mortality cannot be ruled out as an alternative hypothesis.

In addition, the lower rates of mortality in ASW compared to pond water sources suggest lab studies of Bd infection are likely to consistently overestimate transmission rates relative to field or mesocosm studies. More often than not, lab studies use a medium, like ASW, distilled water, or sterilized pond water, to expose hosts to Bd, and these media lack natural Bd predators and competitors, likely leading to low Bd mortality and higher infection rates in hosts compared to field or mesocosm studies, which include Bd predators and competitors (Sauer et al. 2020). Experimental designs for studies that seek to predict Bd transmission should consider including a water source that contains natural and living communities of zooplankton, protozoa, and other microbes, given the influence of pond water sources on Bd zoospore mortality rates found in the present study.

Differences in Bd mortality across water treatments indirectly resulted in differences in infection prevalence with the greatest rates of infection prevalence in ASW compared to all pond water sources. Across treatments, infection prevalence was positively predicted by the concentration of live Bd zoospores. We suggest that consumption of live Bd zoospores by protozoa and zooplankton in pond water could have led to decreases in rates of Bd prevalence in pond water treatments; however, future experiments should test this explicitly. We found no evidence that differences in per capita Bd transmission rates across water treatments directly influenced infection prevalence. Susceptibility of the focal amphibian species and infectivity of Bd zoospores possibly did not vary with exposure to different water sources.

Finally, we found that as a result of similarities in the decomposition rates of Bd zoospores across water sources, concentrations of dead Bd through time did not vary among water sources. These results could suggest that exposure of amphibian hosts to dead Bd and its metabolites, which can induce acquired resistance in amphibian hosts (Ellison et al. 2014, McMahon et al. 2014, Savage et al. 2016), might not commonly vary among local sites, assuming a similar degree of similarity in Bd scavenger and decomposer communities and water quality parameters compared to those in the current study. While we did not directly examine rates of decomposition of Bd metabolites, we might assume similarities in the rates of decomposition of Bd zoospores and metabolites, though future studies should be completed to evaluate this assumption. We know of no other study that has directly estimated decomposition rates of any life stage of Bd, which could be an important consideration in understanding spatial and temporal variability in acquired resistance in natural host populations. Scientists are currently considering how to aid amphibian hosts in developing acquired resistance against Bd as a management strategy to limit the spread of chytridiomycosis. These ongoing efforts could be informed by a better understanding the decomposition of Bd and its metabolites under natural conditions.

While considerable efforts have been undertaken by researchers over the last twenty years to uncover the environmental drivers of the spread of Bd (Kilpatrick et al. 2010, James et al. 2015, Fisher and Garner 2020), additional insights into variability in Bd zoospore mortality, decomposition, and per capita transmission could aid in understanding outbreak heterogeneity among host populations. Our results show that differences in infection prevalence can be mechanistically driven by differences in Bd zoospore mortality but not per capita transmission rates. These results indicate that differences in mortality of Bd zoospores among neighboring sites have the potential to drive local scale patterns of disease dynamics. Further, we show that Bd decomposition is similar across water sources from neighboring sites; these results suggests that exposure of hosts to dead Bd and/or its metabolites, which may lead to acquired resistance, could be similar at small spatial scales. Increased attention towards the effects of local scale variability in pathogen mortality, decomposition, and per capita transmission may increase the efficacy of efforts to surveil and manage wildlife infectious diseases (Cunningham et al. 2017, Barnett and Civitello 2020).

## Author Contributions

S.L.R, T.A.M., D.J.C., and J.R.R designed the experiment; S.A.R. and S.L.R conducted the experiments; T.A.M. completed the qPCR analyses; S.L.R. and D.J.C. analyzed the data; S.L.R. wrote the manuscript; T.A.M, D.J.C, and J.R.R. provided funding, and all authors contributed to editing the manuscript.

## Data Availability

Data will be made available via figshare upon acceptance of the publication.

## Acknowledgements

— This research was supported by grants from the National Science Foundation (J.R.R.: DEB-2017785, DEB-1518681, IOS-1754868, T.A.M.: IOS-1754862, D.J.C: IOS-1755002), National Institutes of Health (J.R.R.: R01TW010286; T.A.M.: 1R01GM135935-01, subaward: KK2022), US Department of Agriculture (2021-38420-34065) to JRR, and the John Wesley Powell Center for Analysis and Synthesis, funded by the U.S. Geological Survey, as a part of the Analyses of Contaminant Effects in Freshwater Systems Working Group to J.R.R. and S.L.R. We would like to acknowledge Joyce Longcore for providing the Bd isolate.

